# Potential of *Bacillus subtilis* as a biological control agent against three rot-causing pathogens in seed yams

**DOI:** 10.64898/2026.05.05.722924

**Authors:** Gideon Darko, Gifty Tetteh, Kelvin Acheampong, Afua Sarpong Sarkodie, Atta Kwesi Aidoo, Daniel Agbetiameh

## Abstract

Ghana is the largest exporter of yams in West Africa. However, yam production, particularly seed yam production, is constrained by storage rot during the off-season. Farmers seldom use synthetic pesticides to control seed yam rot. However, these are costly and pose adverse health risks to farmers. Biological antagonists offer a sustainable, relatively cost-effective, and safe alternative to synthetic pesticides. Therefore, this study aimed to test the efficacy of *Bacillus subtilis* as an alternative to synthetic pesticides. *Bacillus subtilis* supplied through the biofungicide Serenade ASO (Bayer) was assayed against three storage-rot pathogens: *Lasiodiplodia theobromae, Aspergillus niger*, and *Rhizopus* sp. These pathogens were previously isolated from the tissues of rotten seed yams. The efficacy of the bacterium was tested at concentrations of 17 %, 33 %, and 50 % in both in vitro and in vivo bioassays. *B. subtilis* (50 %) completely inhibited the growth (100 %) of *L. theobromae* in the in vitro studies. In contrast, there was little to no growth inhibition of the other two test fungi. In the in vivo assay, *B. subtilis* (50%) significantly (P < 0.01) inhibited *L. theobromae*, resulting in minimal rot lesions. However, *B. subtilis* (50 %) was ineffective against the other two test pathogens, resulting in large rot lesions on the seed yams. This suggests that *B. subtilis* could be an ideal alternative to synthetic pesticides for controlling *L. theobromae* on seed yams.

## Introduction

Yams (*Dioscorea spp*.) constitute a major staple food for about a quarter of the people in Africa (Obidiegwu *et al*., 2020). The crop is cultivated for its edible tubers through vegetative propagation (i.e., minisett and seed yams) across multiple species. The estimated potential yield for yams in Ghana is 52.0 MT/ha. However, the actual yield produced is 18.58 MT/ha (FAOSTAT, 2025; MoFA, 2022). This translates to an unachieved yield gap of 66.6 %. This high yield gap results from several constraints, including a lack of quality planting material, low soil fertility, and pest and disease problems (Osei *et al*., 2019).

Quality planting materials for yams emanate from healthy seed yams. This is a small, whole tuber or cut pieces of yam tubers used as planting material, generally saved from the previous harvest. To obtain healthy seed yams, producers often identify, tag, monitor, and harvest tubers from symptomless mother plants for seed production; a technique described as the Positive Selection (PS) strategy (Kakuhenzire *et al*., 2012). Besides, modern techniques such as tissue culture and aeroponics systems are also being explored for yam propagation (Balogun and Maroya, 2014). However, these latter techniques are not without limitations. The drawbacks associated with these techniques include rainfall, heat shock, soil pests and diseases, and planting time, all of which influence the establishment of tissue culture and aeroponics seedlings in the field (Akom *et al*., 2020).

Seed yams (like ware yams) are highly vulnerable to post-harvest rots caused by fungi, nematodes, and bacteria (Dania *et al*., 2016). Several fungal species, including *Lasiodiplodia theobromae*, have been implicated in causing rots of yam tubers in Ghana (Aidoo, 2015). Similarly, the yam nematode, *Scutellonema bradys*, and root-knot nematode, *Meloidogyne* spp., are well known to cause damage to yams and seed yams during storage in the field after harvest (Sikora *et al*., 2021; Adegbite *et al*., 2005).

While crop infection by these pathogens significantly contributes to the low yields producers attain, their presence during the harvest and postharvest stages can exacerbate the prevailing high yield gap. Timely and effective management of these pathogenic agents is therefore essential during the entire crop value chain. Over the years, the primary strategies for controlling fungal diseases in seed yams have been the use of synthetic fungicides and wax coating (Ntui *et al*., 2021). Recently, the effectiveness of Bordeaux mixture and Carbendazim in managing yam anthracnose and leaf spot disease in seed yams in Ghana was demonstrated (Aidoo, 2024). Regardless of their efficacy, the use of these synthetic fungicides raises important ecological, toxicological risks and safety concerns (Garinie *et al*., 2024; Hashim *et al*., 2023).

Considering the contribution of yams to food security in Africa, it cannot be far-fetched to explore alternative means of managing yam diseases. The potential for biological control through antagonistic microorganisms has been investigated (Lahlali *et al*., 2022; Morales-Cedeño *et al*., 2021). Many biocontrol products involve the deployment of fungi and/or bacteria as active ingredients (Droby *et al*., 2016). Some *Bacillus* species, including *B. subtilis, B. licheniformis, B. pumilis, B. amyloliquifasciens, B. cereus, and B. mycoides*, have been used to inhibit the growth of numerous fungi, including *Phytophthora, Verticillium, Fusarium, Rhizoctonia, and Sclerotinia* species (Dania *et al*., 2016). *Bacillus subtilis*, for instance, exhibits direct and indirect biocontrol strategies to prevent fungal diseases. As part of the direct process, the plant produces a variety of secondary metabolites, hormones, enzymes that break down cell walls, and antioxidants to help protect itself from pest attacks. The application of *Bacillus subtilis* as a biocontrol agent to prevent post-harvest rot in ware yams has been investigated and proven effective against some rot pathogens (Zhou *et al*., 2024; Dania *et al*., 2016). There is, however, little knowledge regarding the viability of employing *Bacillus subtilis* to control postharvest rot of seed yams. Therefore, this study investigated the potency of *Bacillus subtilis* in reducing postharvest storage rot of seed yams.

## Materials and Methods

### Sampling of rotten seed yams

Seed yams of the white yam variety, Dente, showing soft rot symptoms, were collected from yam barns at CSIR-Crops Research Institute, Fumesua, Kumasi, Ghana, and transported to the Pathology laboratory of the same institute for further analysis.

### Isolation and identification of rot pathogens

Infected tissues were excised from the peripheries of the rotten seed yams and surface sterilized in 5 % NaOCl solution for 3 minutes. The excised tissues were washed with three exchanges of deionized water and air-dried under a sterile Laminar flow chamber. The dried diseased tissues were carefully placed on a sterile PDA plate. Petri plates were sealed with parafilm and labelled. The plates were incubated at 28 °C for 3 days. Fungal colonies were sub-cultured onto fresh PDA until pure cultures were obtained. Characteristics of fungal isolates from rotten yam tubers, such as colony colour, colony texture, and spore shapes, were observed using the compound microscope and documented following the protocols described by Mathur and Kongsdal (2003).

### Pathogenicity tests

Inocula from seven-day-old pure cultures of the fungal isolates obtained from rotten seed yam tubers were used for the pathogenicity studies. Seemingly healthy *Dente* seed yam tubers (averaging 800 g) were inoculated with pure cultures of the fungal isolates. A sterile cork-borer (dia. 5 mm) was used to cut plugs from the 7-day-old cultures of the fungal isolates. This was placed in 1 cm-deep holes created in the seed yam tubers and sealed with melted candle wax. Each fungal isolate was replicated 3 times at 3 different positions, 10 cm apart, on seed yam tubers. Controls were set up in which no fungal organism was inoculated into the holes created in the seed yams. Re-isolations of the rot pathogens were conducted, and their characteristics were compared with the previous ones.

### Efficacy test of *Bacillus subtilis*

#### Preparation of test concentrations

*Bacillus subtilis* was applied via the commercial bio-pesticide, Serenade ASO, which contains *B. subtilis* strain QST 713 as the active microbial agent. Three concentrations of the active microbial agent were used to represent the treatments, in accordance with the manufacturer’s recommendations. Additionally, Bordeaux mixture (Caldo Bordeles) and water were used as positive and negative controls, respectively (Table 1).

**Table 1.** Treatment concentrations of the test product and controls tested against rot pathogens.

| Treatment | Amount of Product | Amount of Solvent (Water) |
| --- | --- | --- |
| 100 ml of Serenade ASO | 17 ml | 83 ml |
| 100 ml of Serenade ASO | 33 ml | 67 ml |
| 100 ml of Serenade ASO | 50 ml | 50 ml |
| 200 ml of Caldo Bordeles | 2 g | 200 ml |
| 100 ml of Water | - | 100 ml |

#### Anti-fungal activity of *Bacillus subtilis* on seed yam rot pathogens

Three test fungi, *Lasiodiplodia theobromae, Aspergillus niger*, and *Rhizopus* sp., isolated and identities confirmed previously, were used in this experiment. The poisoned food technique was used to investigate the efficacy of *Bacillus subtilis*. Potato Dextrose Agar (PDA) was amended with 1 ml of the three concentrations of Serenade ASO described previously (Table 1). The amended medium was inoculated centrally with 5-mm-diameter discs of 7-day-old cultures of the test fungi. Three replications were set for each treatment. Controls were set up in which a PDA with distilled water was inoculated with test fungi. Further, controls were set up in which a solution of Bordeaux mixture was added to PDA in Petri dishes. The whole setup was incubated at 28 °C. The radius of the growing fungal colonies was measured and recorded at 24 h intervals for four consecutive times using a ruler.

Data collected was used to calculate the Mycelial Growth Inhibition (MGI) for each Serenade ASO concentration and Bordeaux mixture using the following formulae proposed by (Wonglom *et al*., 2019):

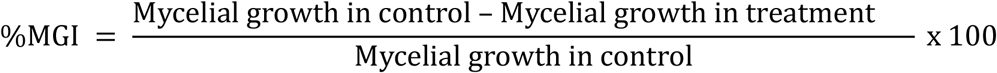

#### Data analysis

The results were analyzed using analysis of variance (ANOVA) in the statistical software Statistix (10^th^ edition). Treatment means were separated using Tukey’s Honestly Significant Difference at 1 % significance level.

## Results

### Identification of fungal isolates and pathogenicity studies

Three fungal species, *viz*., *Lasiodiplodia theobromae, Aspergillus niger, and Rhizopus sp*., *were identified based on* morphological characteristics (Figure 1). The fungal species *L. theobromae* showed a dark grey colony colour, a cottony texture and ellipsoid spores. *Aspergillus niger*, on the other hand, had a dark colony colour, which was powdery and spherical conidia. *Rhizopus* spp. expressed a dark colony colour, cottony, and oval-shaped sporangiospores (Table 2).

**Table 2.** Characteristics of isolated fungal pathogens.

| Fungal Pathogens | Colony colour | Colony texture | Spore shape |
| --- | --- | --- | --- |
| *L. theobromae* | Dark grey | Cottony | Ellipsoid |
| *A. niger* | Black | Powdery | Spherical |
| *Rhizopus* sp. | Grey | Cottony | Oval |

**Figure 1.**
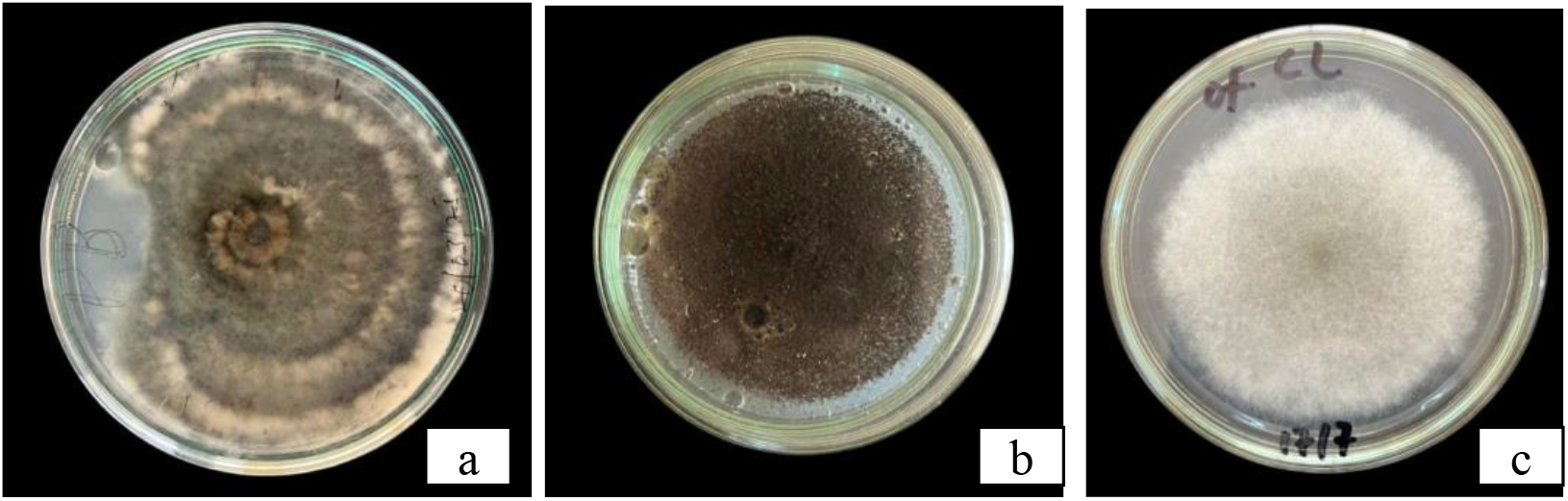
Morphological features of isolated fungal pathogens from rotten seed yam tubers: (a) culture plate of *L. theobromae*, (b) culture plate of *A. niger*, (c) culture plate of *Rhizopus* sp.

Each fungal species caused dry rot lesions except *Rhizopus* sp., which caused soft rot when inoculated into healthy yam tubers (Figure 2).

**Figure 2.**
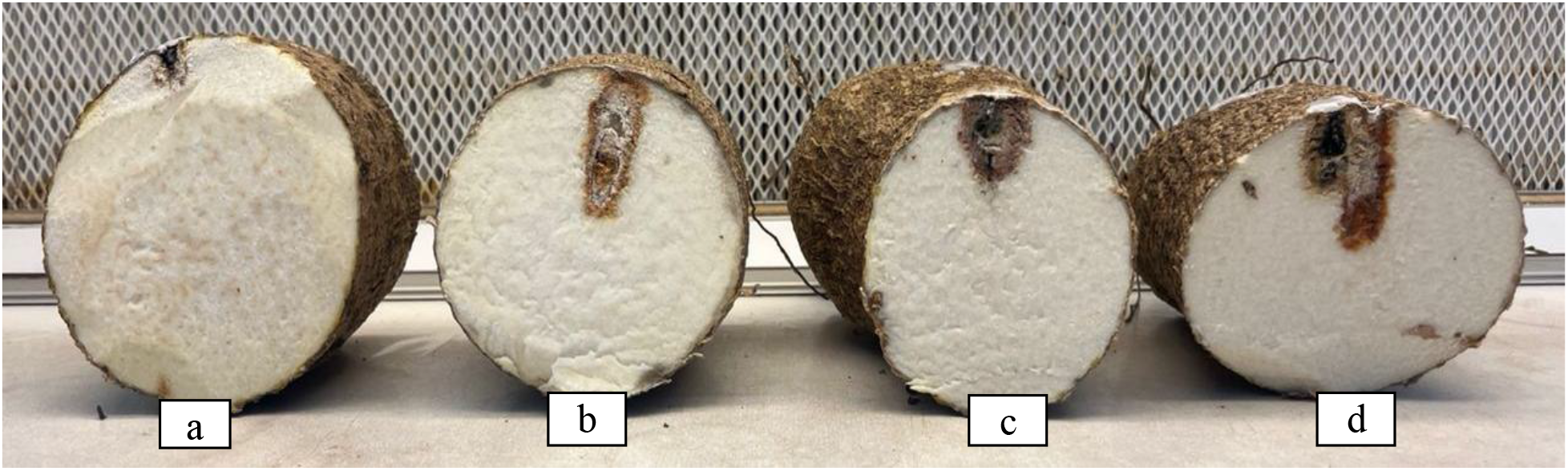
Cross-section of seed yam showing lesions from pathogenicity tests (a) agar block, (b) *L. theobromae*, (c) *Aspergillus niger*, (d) *Rhizopus* sp.

### Effects of *Bacillus subtilis* against isolated seed yam rot fungi in vitro

Mycelial growth inhibition by *Bacillus subtilis* was more significant in *Lasiodiplodia theobromae* compared to *Aspergillus niger* and *Rhizopus* sp. While no inhibition was observed for *L. theobromae* at 17 %, 33 %, or 50 % concentrations after 24 hours of incubation, substantial inhibitory effects emerged after 96 hours, with percent inhibition increasing in a concentration-dependent manner. In contrast, Bordeaux mixture exhibited comparatively lower efficacy against *L. theobromae* after the same incubation period.

For *A. niger*, no inhibition was observed at 24 hours across all concentrations of *B. subtilis*. By 96 hours, minimal inhibition was recorded only at the 33% and 50% concentrations, with no significant differences compared to control treatments.

*Rhizopus* sp. displayed complete resistance to *B. subtilis* treatment, with no observable inhibition at any concentration or time point, consistent with control values.

**Table 3.** Inhibitory effect of *Bacillus subtilis* against isolated seed yam fungal rot pathogens in an *in vitro* assay.

| Fungal rot<br>pathogens | Concentration of *B. subtilis* |  |  |  |  |  | Bordeaux<br>mixture |  | Control |  |
| --- | --- | --- | --- | --- | --- | --- | --- | --- | --- | --- |
|  | 17 ml |  | 33 ml |  | 50 ml |  | 200 ml |  | 100 ml |  |
|  | MGR | MGI | MGR | MGI | MGR | MGI | MGR | MGI | MGR | MGI |
| *L. theobromae* | 0.14 | 66.0 | 0.10 | 76.3 | 0.0 | 100 | 0.3 | 29.2 | 0.4 | 0.0 |
| *A. niger* | 0.42 | 0.0 | 0.40 | 5.0 | 0.30 | 34.4 | 0.3 | 0.0 | 0.4 | 0.0 |
| *Rhizopus* spp. | 0.42 | 0.0 | 0.42 | 0.0 | 0.42 | 0.0 | 0.3 | 0.0 | 0.4 | 0.0 |
| HSD (0.01) | 0.07 | 17.24 | 0.17 | 40.35 | 0.21 | 49.36 | 0.22 | 53.4 | 0.0 | 0.0 |
| CV (%) | 6.04 | 21.23 | 15.11 | 40.69 | 24.42 | 30.1 | 16.3 | 151.3 | 0.0 | 0.0 |

The data represent the mean values of three replicates.

### Effects of *Bacillus subtilis* against isolated seed yam rot fungi in vivo

*Bacillus subtilis* at a concentration of 50 % was tested against the isolated pathogens in vivo on healthy seed yams. *B. subtilis*, when simultaneously inoculated with test fungi, showed rot lesions in the seed yams (Figure 3). Seed yam inoculated with the bacterium and *L. theobromae* showed minimal rot lesions compared to lesions created by the other two rot fungi inoculated with *B. subtilis*.

**Figure 3.**
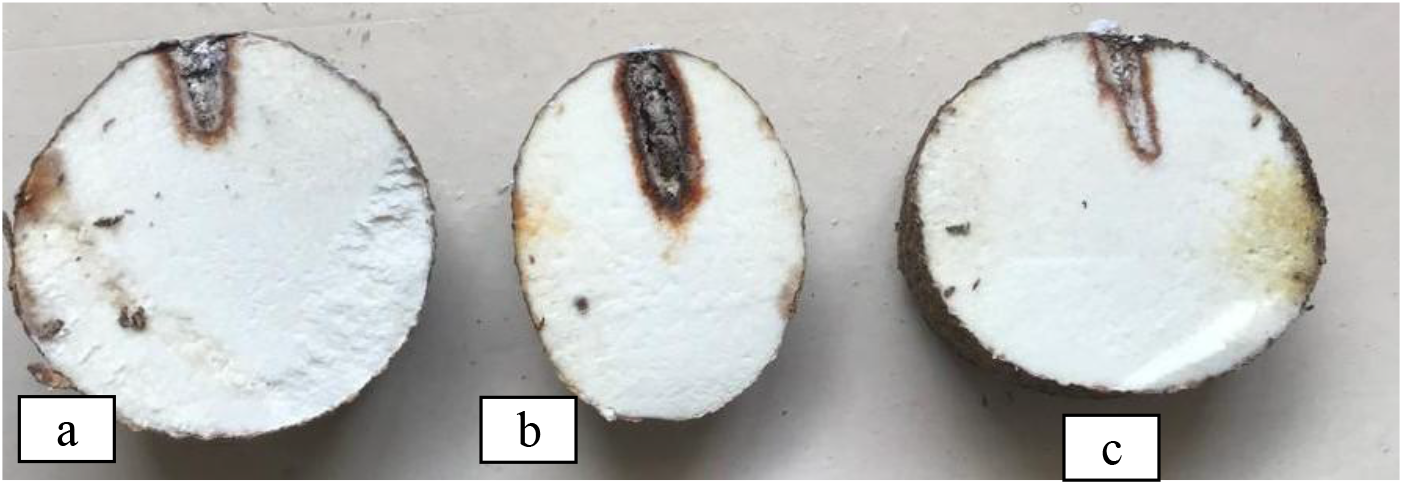
In vivo assay of *B. subtilis* against isolated fungal pathogens (a) *B. subtilis and L. theobromae*, (b) *B. subtilis and A. niger*, (c) *B. subtilis and Rhizopus sp*.

## Discussion

Seed yam rot disease is associated with the post-harvest handling of yam tubers. Some of the major microorganisms causing rot diseases in seed yams include *Aspergillus flavus, Aspergillus niger, Lasiodiplodia theobromae, Fusarium oxysporum, Fusarium solani, Penicillium chrysogenum, Penicillium oxalicum, Trichoderma viride, Rhizoctonia* spp., and *Rhizopus nodosus* (Okigbo, 2004; Aidoo, 2011). Three of the fungi, *L. theobromae, A. niger*, and *Rhizopus* sp., were isolated and identified as rot-causing organisms in this study. Little work has been done to investigate the use of biological agents as biopesticides to control seed yam post-harvest storage rot; hence, this study explored the potential of *Bacillus subtilis*.

Previous studies have highlighted the antagonistic properties of *Bacillus subtilis* against various postharvest spoilage fungi in ware yams. *B. subtilis* exhibited significant antagonism against *L. theobromae* and *A. niger*, with 52.5 % and 60 % mycelial growth inhibition, respectively (Okigbo, 2004). These findings align with the in vitro efficacy of *B. subtilis* reported in this study. At a 50 % concentration, *B. subtilis* completely (100 %) inhibited the mycelial growth of *L. theobromae*. However, the discrepancy in the efficacy of *B. subtilis* against *A. niger* between this study and Okigbo (2004) could be attributed to differences in experimental conditions, such as B. subtilis concentration or strain.

The work of Okigbo (2004) further supports the use of *B. subtilis* as an effective biocontrol agent against yam rot pathogens. That study noted that yams inoculated with *B. subtilis* did not develop rot, whereas those inoculated with *A. niger* and *L. theobromae* showed considerable rot. Rot developed when *B. subtilis* was inoculated with these fungi simultaneously. This suggests that while *B. subtilis* has antagonistic properties; its efficacy can be compromised under certain conditions. This finding is consistent with the reduced *in vivo* effectiveness of *B. subtilis* demonstrated in this study. Despite its effective *in vitro* performance, *B. subtilis* could not completely prevent rot caused by *L. theobromae in vivo*.

The observed limitations of *B. subtilis* in controlling *A. niger* and *Rhizopus* sp. in vivo also echo the challenges highlighted by the study by Okigbo (2004). He stated that ‘while *B. subtilis* is effective in laboratory settings, its real-world application is complicated by factors such as the bacterium’s ability to penetrate through wounds in yam tubers caused by insects, nematodes, and poor handling practices’. These wounds provide entry points for fungi, which thrive under favourable environmental conditions, such as high moisture levels, optimal temperature, and pH. The challenge of maintaining effective concentrations of *B. subtilis* in such complex environments may explain its reduced performance in our in vivo assays.

Moreover, *B. subtilis* is not known to cause rot in yams or seed yams and is considered a beneficial soil bacterium with antifungal properties (Okigbo, 2004). The specific interactions between *B. subtilis* and the various fungal pathogens in this study highlight the complexity of biological control. The ineffectiveness of *B. subtilis* against *A. niger* and *Rhizopus* sp. in our study, despite its known broad-spectrum antifungal activity, suggests that these pathogens may possess resistance mechanisms or that the biocontrol conditions were suboptimal for *B. subtilis* to exert its activity.

The potential of *Bacillus subtilis* as a biocontrol agent for managing storage rot in seed yams has been explored, highlighting its efficacy against *Lasiodiplodia theobromae* while revealing limitations in controlling *Aspergillus niger* and *Rhizopus* sp., particularly under in vivo conditions. *B. subtilis* demonstrated complete inhibition of *L. theobromae* in vitro, yet its performance against other pathogens was less effective, suggesting that further optimization is needed.

It is recommended that future efforts should focus on optimizing application methods and exploring synergistic use with other biocontrol agents and other management practices to offer more comprehensive protection against a broad spectrum of rot pathogens.

## Acknowledgments

We thank the Plant Pathology Division of CSIR-Crops Research Institute, Fumesua, Kumasi, Ghana, for allowing us to use their laboratory for this study.

## References

Adegbite, A.A., Adesiyan, S.O., Agbaje, G.O. and Omoloye, A.A. (2005) “Host suitability of crops under Yam intercrop to root-knot nematode (Meloidogyne incognita Race 2) in South-Western Nigeria,” Journal of Agriculture and Rural Development in the Tropics and Subtropics (JARTS), 106(2), pp. 113–118.

Aidoo, A.K. (2011). Yam tuber rot: Identification and control of pathogens in storage. PhD Thesis. Available at: https://ir.knust.edu.gh/handle/123456789/273 (Accessed: March 27, 2024).

Aidoo, A.K. (2015) “Improving pit storage systems to reduce rots of whiteyam (Dioscorea rotundata) in Ghana,” Science Journal of Agricultural Research and Management, 2015. (Accessed: February 5, 2024).

Akom, M., Ennin, S.A., Osei, K., Appiah-Kubi, D. and Quain, M.D. (2020) “Field performance of seed yam (Dioscorea rotundata Poir) derived from tissue culture and aeroponics seedlings,” EC Agriculture, 6, pp. 67–75.

Balogun, M.O. and Maroya, N. (2014) “Status and prospects for improving yam seed systems using temporary immersion bioreactors,” African Journal of Biotechnology, 13(15). Available at: 10.5897/AJBX2013.13522.

Dania, V.O., Fadina, O.O., Ayodele, M. and Kumar, P.L. (2016) “Evaluation of isolates of Trichoderma, Pseudomonas and Bacillus species as treatment for the control of post-harvest fungal rot disease of yam (Dioscorea spp.),” Archives of Phytopathology and Plant Protection, 49(17– 18), pp. 456–470. Available at: 10.1080/03235408.2016.1231496.

Droby, S., Wisniewski, M., Teixidó, N., Spadaro, D. and Jijakli, M.H. (2016) “The science, development, and commercialization of postharvest biocontrol products,” Postharvest Biology and Technology, 122, pp. 22–29.

Garinie, T., Nusillard, W., Lelièvre, Y., Taranu, Z.E., Goubault, M., Thiéry, D., Moreaua, J. and Louâprea, P. (2024) “Adverse effects of the Bordeaux mixture copper-based fungicide on the non-target.” Pest Management Science 1–10 (Accessed: October 21, 2025).

Hashim, M., Al-Attar, A.M. and Zeid, I.M.A. (2023) “Toxicological effects of carbendazim: a review,” Acta Sci. Med. Sci, 7. (Accessed: October 21, 2025).

Kakuhenzire, R., Lemaga, B., Tibanyendera, D., Borus, D., Kashaija, I., Namugga, P. and Schulte-Geldermann, E. (2012) “Positive selection: A simple technique for improving seed potato quality and potato productivity among smallholder farmers,” in II All Africa Horticulture Congress 1007, pp. 225–233. (Accessed: May 15, 2024).

Lahlali, R., Ezrari, S., Radouane, N., Kenfaoui, J., Esmaeel, Q., El Hamss, H., Belabess, Z. and Barka, E.A. (2022) “Biological Control of Plant Pathogens: A Global Perspective,” Microorganisms, 10(3), p. 596. Available at: 10.3390/microorganisms10030596.

Mathur, S.B. and Kongsdal, O. (2003) “Common laboratory seed health testing methods for detecting fungi, 2nd Edition. International Seed Testing Association. Switzerland.”

Ntui, V.O., Uyoh, E.A., Ita, E.E., Markson, A.-A.A., Tripathi, J.N., Okon, N.I., Akpan, M.O., Phillip, J.O., Brisibe, E.A., Ene-Obong, E.-O.E. and Tripathi, L. (2021) “Strategies to combat the problem of yam anthracnose disease: Status and prospects,” Molecular Plant Pathology, 22(10), pp. 1302–1314. Available at: 10.1111/mpp.13107.

Obidiegwu, J.E., Lyons, J.B. and Chilaka, C.A. (2020) “The Dioscorea Genus (Yam)—An Appraisal of Nutritional and Therapeutic Potentials,” Foods, 9(9), p. 1304. Available at: 10.3390/foods9091304.

Okigbo, R.N. (2004) “A review of biological control methods for postharvest yams (Dioscorea spp.) in storage in South Eastern Nigeria,” KMITL Sci J, 4(1), pp. 207–215.

Osei, K., Ennin, S., Aighewi, B., Aidoo, A., Lamptey, J., Mochiah, M., Aihebhoria, D., Adomako, J., Appiah-Kubi, Z., Godfried, O.-M., Asante, B., Adu, J. and Osuman, A. (2019) “Enhancing Productivity of Farmer-Saved Seed Yam in Ghana,” African Crop Science Journal, 27(4), pp. 631– 640. Available at: 10.4314/acsj.v27i4.6.

Sikora, R.A., Desaeger, J. and Molendijk, L. (eds.) (2021) Integrated nematode management: state-of-the-art and visions for the future. 1st ed. UK: CABI. Available at: 10.1079/9781789247541.0000.

Wonglom, P., Daengsuwan, W., Ito, S. and Sunpapao, A. (2019) “Biological control of Sclerotium fruit rot of snake fruit and stem rot of lettuce by Trichoderma sp. T76-12/2 and the mechanisms involved,” Physiological and Molecular Plant Pathology, 107, pp. 1–7.

Zhou, J., Chang, X., Liu, J., Yang, X., Zhang, H. and Chen, X. (2024) “Study on The Effect of Bacillus Subtilis on The Fungal Community of Fresh Cut Yam,” in BIO Web of Conferences. EDP Sciences, p. 03021. (Accessed: October 21, 2025).

